# Programmed hierarchical patterning of bacterial populations

**DOI:** 10.1101/204925

**Authors:** Christian R. Boehm, Paul K. Grant, Jim Haseloff

## Abstract

Modern genetic tools allow the dissection and emulation of fundamental mechanisms shaping morphogenesis in multicellular organisms. Several synthetic genetic circuits for control of multicellular patterning have been reported to date. However, hierarchical induction of gene expression domains has received little attention from synthetic biologists, despite its importance in biological self-organization. We report the first synthetic genetic system implementing population-based AND logic for programmed autonomous induction of bacterial gene expression domains. We develop a ratiometric assay for bacteriophage T7 RNA polymerase activity and use it to systematically characterize different intact and split enzyme variants. We then utilize the best-performing variant to build a three-color patterning system responsive to two different homoserine lactones. We validate the AND gate-like behavior of this system both in cell suspension and in surface culture. Finally, we use the synthetic circuit in a membrane-based spatial assay to demonstrate programmed hierarchical patterning of gene expression across bacterial populations.

**Abbreviations:**

3OC6HSL: N-(3-oxohexanoyl)-L-homoserine lactone
3OC12HSL: N-(3-oxododecanoyl)-L-homoserine lactone
CFP: cyan fluorescent protein
IPTG: isopropyl-β-D-thiogalactopyranoside
Parts Registry: MIT Registry of Standard Biological Parts
PI: positional information
RD: reaction-diffusion
RFP: red fluorescent protein
RFU: relative fluorescence units
s.d.: standard deviation
T7RNAP: bacteriophage T7 RNA polymerase
YFP: yellow fluorescent protein

## Introduction

The hierarchical organization of multicellular organisms builds on mechanical and chemical interactions. Cells sense those cues in their environment and accordingly modulate their own metabolism as well as their intercellular communication with neighbours. At the population level, the spatiotemporal interplay of multiple processes leads to the emergence of self-organization through mechanisms such as symmetry breaking, domain induction, and boundary formation^1^. Despite great advances made in the biological sciences over the past decades, many of the complex mechanisms underlying morphogenesis remain elusive.

In efforts to explain multicellular patterning, two types of models have been preeminent to date: the reaction-diffusion (RD) model proposed by Alan Turing^2^, and the positional information (PI) model (also known as the “French flag model”) originating from Lewis Wolpert^3^. RD-type systems are characterized by self-organized spatial patterns emerging from interacting positive- and negative-feedback loops in response to two diffusible morphogens. By contrast, PI-type models employ a single predefined morphogen gradient, which is interpreted by receiving cells according to the local concentration of the morphogen. Though the two models are conceptually distinct, they are not necessarily mutually exclusive^4^.

Empowered by modern genetic techniques, molecular biologists are now striving to not only dissect developmental processes, but to exploit their modularity for the design of custom living systems^5^. Biological self-organization is a powerful tool for bioprocessing and remediation in tailored microbial consortia, for sustainable bioproduction in novel plant compartments, or for applications in regenerative medicine enabled by engineered vertebrate tissues. A fundamental requirement for harnessing this potential is control over the differentiation of cell types to create domains of gene expression in spatially organized patterns. To date, several synthetic biological circuits capable of multicellular patterning have been reported, predominantly implemented either by RD-type^6,7^ or by PI-type^8–11^ mechanisms.

However, mechanisms for hierarchical patterning have received little attention from synthetic biologists to date. In various examples of morphogenesis, such as vulval induction in nematodes^12^, the ABC model of flower development^13^, or mesoderm induction in vertebrates^14^, we can observe nested domains of gene expression. To emulate biological self-organization on this level of complexity, we require control over hierarchical induction of new domains within existing patterns.

Approaching this challenge, we report the implementation of a synthetic genetic circuit that controls emergence of a new domain of gene expression at the interface of existing bacterial populations. To best of our knowledge, this is the first description of a synthetic genetic circuit implementing AND logic for autonomous hierarchical patterning at the population scale. A previously reported population-based edge detector has implemented AND (NOT (NOT)) logic in response to light projected through a mask^15^. By contrast, the circuit introduced here is designed to establish two layers of patterning in the absence of an externally defined spatial input.

As intercellular signals mediating domain induction, we utilize the diffusible small molecule signals N-(3-oxohexanoyl)-L-homoserine lactone (3OC6HSL) and N-(3-oxododecanoyl)-L-homoserine lactone (3OC12HSL). These compounds are derived from the bacterial quorum sensing systems from *Vibrio fischeri*^16^ and *Pseudomonas aeruginosa*^17^, respectively. Both systems employ a single biosynthetic enzyme to produce a diffusible signal (LuxI / LasI), and a single receiver protein (LuxR / LasR) to activate transcription of a target gene controlled by their cognate promoters (P_Lux_ / P_Las_) upon signal binding^18^. Both signalling systems have been previously combined as part of synthetic circuits, but by default exhibit significant levels of crosstalk^19–24^. Overcoming this complication, we have recently reported an intercellular signalling system minimizing crosstalk between 3OC6HSL and 3OC12HSL in the same cell^24^.

In this work, we combine the improved intercellular signalling system with the transcriptional output from an orthogonal split bacteriophage T7 RNA polymerase (T7RNAP) ^25–29^ to demonstrate hierarchical induction of gene expression domains at the population scale. A first layer of patterning is established via promoters responding to the two different homoserine lactones by induction of cyan (CFP) and yellow fluorescent protein (YFP), respectively. On top of these domains, a second layer of patterning is generated by expression of red fluorescent protein (RFP) from a T7 promoter. With each of the two homoserine lactones inducing expression of one half of the split T7RNAP protein, its transcriptional activity is limited to where the two diffusible signals coincide. First, we systematically characterize the activity of different *T7RNAP* genes using a ratiometric strategy to identify the split variant of highest dynamic range. Then we utilize this to implement a synthetic three-color AND gate based on split T7RNAP responsive to 3OC6HSL and 3OC12HSL in cell suspension and in surface cultures of *Escherichia coli*. Finally, we demonstrate that the synthetic circuit autonomously mediated programmed emergence of a new gene expression domain at the interface of two bacterial populations (Fig. 1).

**Figure 1.**
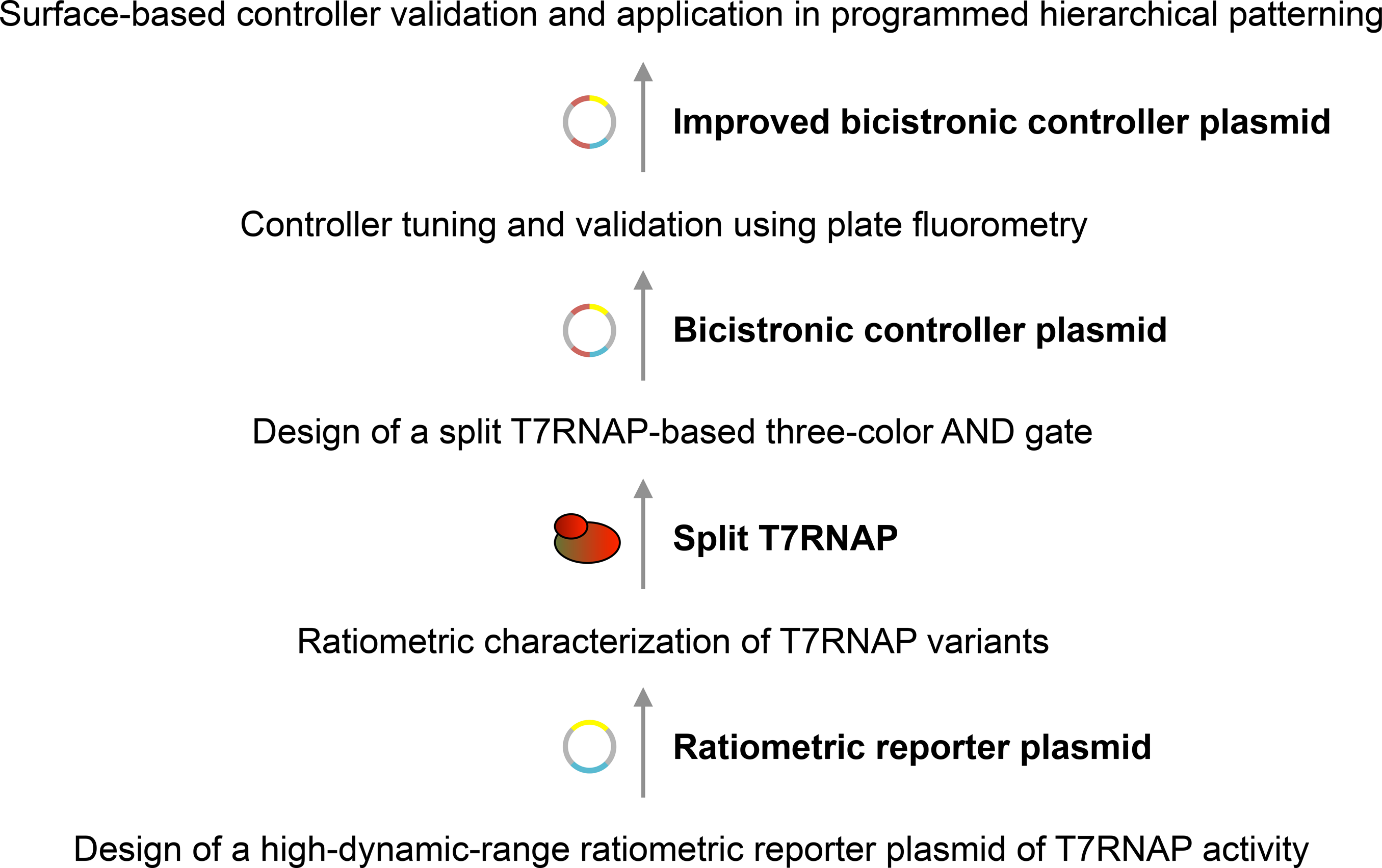
Design workflow of a genetic circuit for synthetic hierarchical patterning.

## Results

### Design of a high-dynamic-range ratiometric reporter plasmid for T7RNAP activity

Rational design of a synthetic circuit generating desired multicellular behavior requires a quantitative understanding of its core genetic components. To increase robustness in measurements of activity from the T7 promoter, we extended a previously reported ratiometric strategy^30,31^ to T7RNAP-driven gene expression: in our ratiometric reporter plasmids, the yellow fluorescent protein variant *mVenus*^32^ served as primary reporter of T7RNAP activity. In addition, the ratiometric reporter plasmids encoded a cyan fluorescent protein variant *mTurquoise2*^33^ which was constitutively expressed under control of reference promoter P_J23101_ and reference RBS_B0034_ (Parts Registry). Relative promoter activity could be expressed as the average YFP over CFP fluorescence intensity per cell during exponential growth phase^30,31^. This indicator was chosen in order to correct for variation in cellular gene expression capacity due to environmental conditions.

We designed a small library of plasmid reporters for T7RNAP activity to identify a combination of promoter, 5’UTR, and copy number capable of supporting high dynamic range in reporter induction. This library embraced all combinations of (i) the wildtype T7 promoter P_T7_^34–36^ or a mutant promoter P_T7_(-3G) (~20% activity of the consensus sequence) ^37^, (ii) the 5’UTR from bacteriophage T7 gene 10 RBS_T7g10_^38^ or a synthetic 5’UTR embracing the reference RBS_B0034_ (Parts Registry), and (iii) the high copy number backbone pSB1A3 (pMB1 ori, Parts Registry) or the low copy number backbone pSB4A5 (pSC101 ori, Parts Registry). Notably, these plasmids included a LacI generator cassette to reduce leaky expression of *T7RNAP* from the genome of *T7 Express E. coli*.

To quantify relative activity from the T7 promoter across the small library of ratiometric reporters outlined above, we introduced the plasmids individually into *T7 Express E. coli* encoding a single genomic copy of *T7RNAP* under control of the *lac* operon. After induction of *T7RNAP* expression in transformed *T7 Express E. coli* using a range of isopropyl-β-D-thiogalactopyranoside (IPTG) concentrations (Supplementary Fig. S1), we performed ratiometric assays in a plate fluorometer to measure CFP and YFP fluorescence intensities and optical density over time for measurement of relative promoter activity (Fig. 2a; see Methods for details). Taking this experimental approach, we found that combining the mutant P_T7_(-3G) promoter with reference RBS_B0034_ on a low copy number backbone produced the highest induction of relative activity from the T7 promoter after addition of inducer (7.58 ± 0.08-fold).

**Figure 2.**
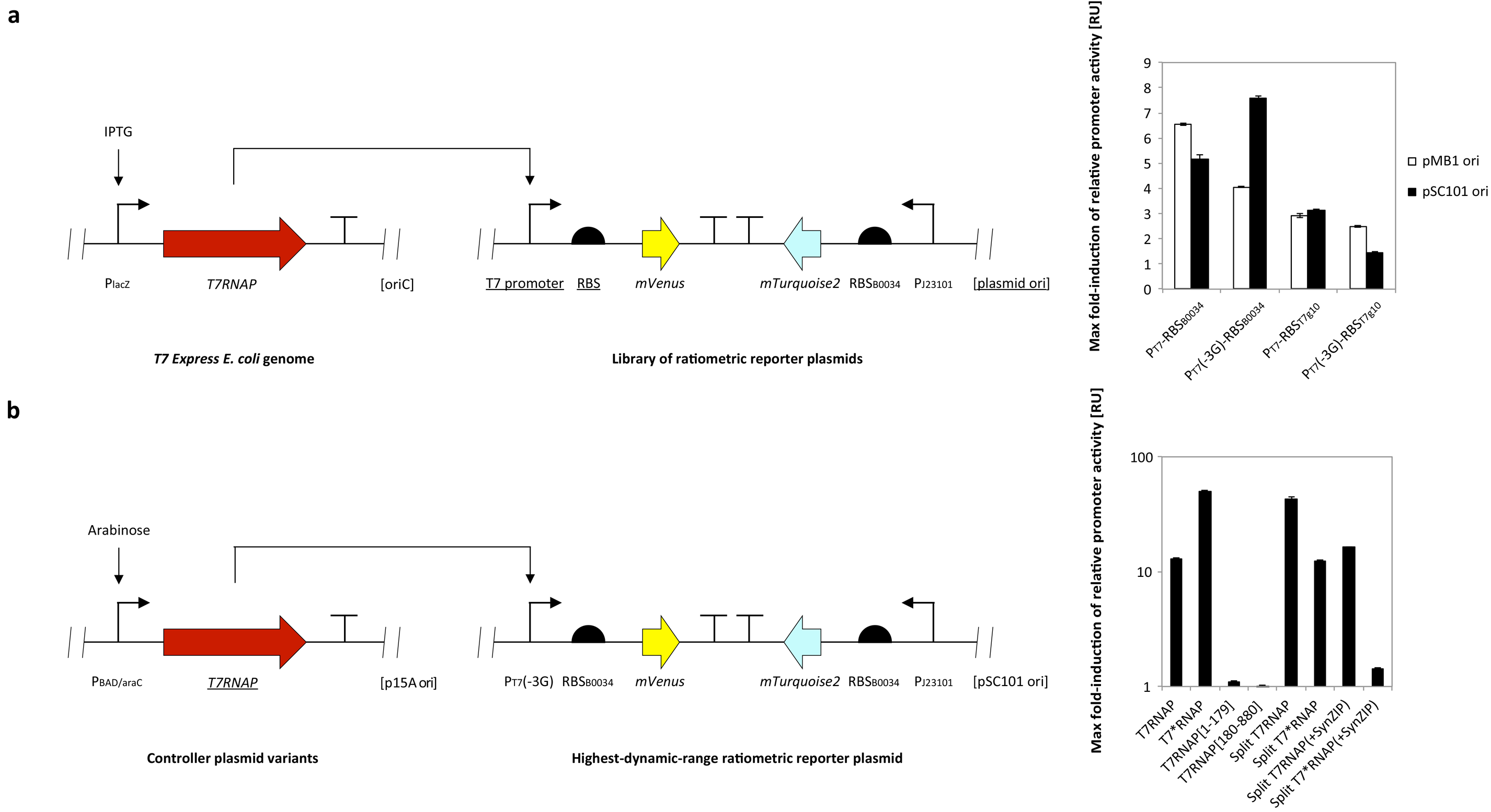
Ratiometric characterization of T7RNAP activity. (**a**) Comparison of different ratiometric reporter plasmids of T7RNAP activity. Ratiometric reporter plasmids combining different variants of the T7 promoter, 5’-UTRs, and origins of replication (underlined) were introduced into *T7 Express E. coli*. The maximum fold-induction of relative activity from the T7 promoter in response to a range of IPTG concentrations (see Supplementary Fig. S1 for induction curves was quantified using a plate fluorometer-based assay (see Methods for details, and is reported relative to the absence of inducer. Error bars represent the s.d. of average values yielded between 3 biological replicate experiments performed on different days. (**b**) Ratiometric characterization of intact and split variants of T7RNAP. *TOP10 E. coli* were co-transformed by the highest dynamic range ratiometric reporter p4g3VT tested under (a and different controller plasmids encoding intact or fragmented variants of the *T7RNAP* gene (underlined) under control of the arabinose-inducible P_BAD/araC_ promoter. The maximum fold-induction of relative activity from the T7 promoter in response to a range of arabinose concentrations was quantified as described above (see Supplementary Fig. S2 for induction curves), and is reported relative to the absence of inducer. Error bars represent the s.d. of average values yielded between 3 biological replicate experiments performed on different days.

### Ratiometric characterization of intact and split T7RNAP variants

Next, we used the reporter plasmid displaying the highest dynamic range in *T7 Express E. coli* for measurement of the activity of different intact and split T7RNAP variants. We cotransformed *TOP10 E. coli* with the ratiometric reporter plasmid p4g3VT and controller plasmids encoding intact and fragmented *T7RNAP* genes under control of the arabinose-inducible P_BAD/araC_ system^26^. The LacI generator was removed from the ratiometric reporter plasmid p4g3VT_LacI_ for this purpose. Performing ratiometric assays with a range of arabinose concentrations in a plate fluorometer-based format, we confirmed that induction of relative activity from the T7 promoter was negligible for both of the individual N-terminal (T7RNAP[1-179]: 1.10 ± 0.01-fold) and C-terminal (T7RNAP[180-880]: 0.94 ± 0.05-fold) fragments of T7RNAP compared to the intact enzyme (T7RNAP: 12.9 ± 0.3-fold; Fig. 2b). High induction of T7RNAP (i.e. at inducer concentrations exceeding 1 mM arabinose) led to a drop in relative promoter activity (see Supplementary Fig. S2). Incorporation of the mutation R632S^39^ into the intact wildtype enzyme increased the maximum induction of relative promoter activity almost by a factor of 4 (T7*RNAP: 50.1 ± 0.9-fold). Furthermore, the inducer concentration required for half maximal activity was increased approximately 30-fold. By contrast to T7RNAP, no inverse correlation between inducer concentration and promoter activity was observed for T7*RNAP up to 50 mM arabinose. Splitting wildtype T7RNAP between residues 179 and 180^25,26^ increased its dynamic range by over a factor of 3 (split T7RNAP: 43 ± 2-fold). In contrast to the intact enzyme, incorporation of the R632S mutation decreased the maximum induction of relative promoter activity in split T7RNAP (split T7*RNAP: 12.3 ± 0.3-fold). Adopting an approach previously taken^27^, we also tested whether the addition of SynZIP protein-protein interaction domains^40,41^ to wildtype and mutant (R632S) variants of split T7RNAP increased their dynamic range. Compared to enzyme modifications described above, application of SynZIP domains further increased the inducer concentration required for half maximal activity (i.e. over 100-fold relative to T7RNAP; see Supplementary Fig. S2). However, neither modified variant exceeded wildtype split T7RNAP in maximum induction of activity from the T7 promoter (split T7RNAP(+SynZIP): 16.3 ± 0.3-fold; split T7*RNAP(+SynZIP): 1.42 ± 0.04-fold). From the ratiometric assays performed, we concluded that split T7RNAP was the most promising nicked variant to employ in our synthetic patterning circuit.

### Design of a split T7RNAP based three color AND gate responsive to homoserine lactones

Having validated the activity of split T7RNAP, we utilized this enzyme to design a three-color synthetic transcriptional AND gate. To this end, we assembled a controller plasmid embracing two bicistronic operons composed of *T7RNAP[1-179]* and *mVenus*, and *T7RNAP[180-880]* and *mTurquoise2*, respectively (Fig. 3). The downstream fluorescent proteins were included to serve as visual indicators of expression of the individual T7RNAP fragments *in vivo*. The two bicistronic operons were controlled by hybrid promoters responsive to either 3OC12HSL (P_Las81*_) or 3OC6HSL (P_Lux76*_), respectively, alongside weak ribosomal binding sites RBS_B0033_ (Parts Registry). The promoters in question were derived from our previously reported system for orthogonal intercellular signalling^24^, and contained mutations shown to reduce basal activity from the Lux promoter^42^. Cognate repressors LuxR and LasR were constitutively expressed from the vector backbone pR33S175. We also constructed a RFP reporter plasmid of T7RNAP activity p4g3R based on the design principles proven earlier (see Fig. 2a): the *mRFP1* gene was controlled by P_T7_(-3G) and RBS_B0034_ on a low copy number pSC101 backbone. By contrast to ratiometric reporter plasmids, p4g3R lacked the P_J23101_-*mTurquoise2* reference operon. In *T7Express E. coli*, p4g3R was validated capable of reporting induction of T7 promoter activity in response to IPTG (Supplementary Fig. S3).

**Figure 3.**
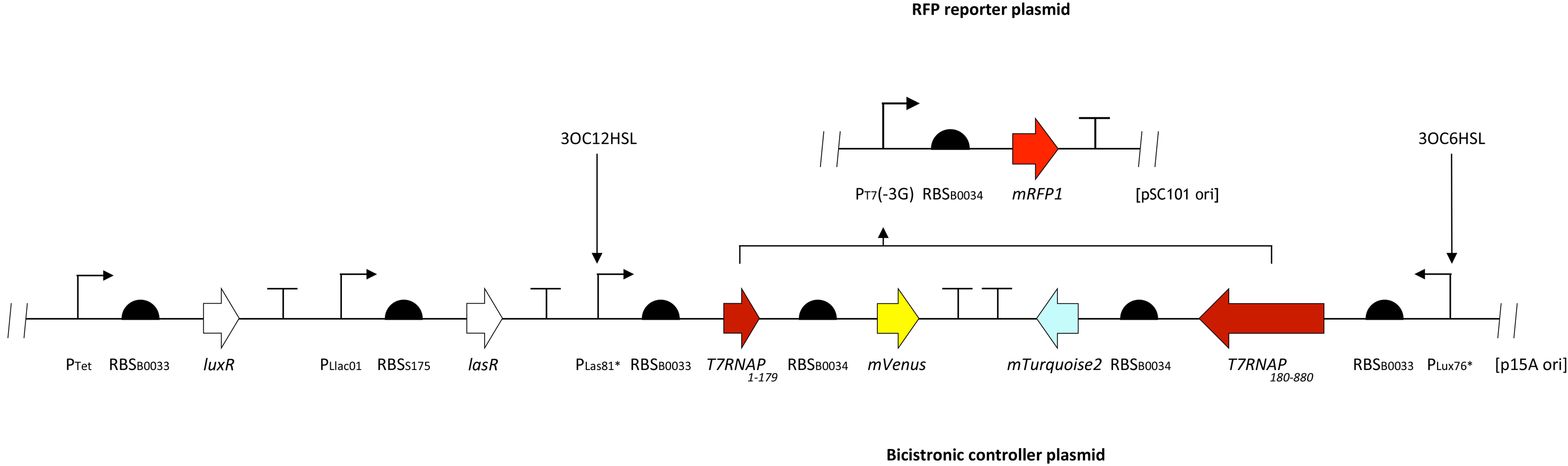
Design of a split T7RNAP-based three-color circuit responsive to homoserine lactones. Shown are schematic representations of the *mRFP1*-expressing reporter of T7RNAP activity p4g3R (see Supplementary Fig. S3 for induction curve) and the bicistronic controller plasmid pCRB DRT7VTPLux*500 encoding two bicistronic operons responding to 3OC6HSL and 3OC12HSL by induction of *T7RNAP[180-880]* and *mTurquoise2*, or *T7RNAP[1-179]* and *mVenus*, respectively.

First, we tested the behavior of the two individual controller half circuits. For this purpose, half-circuit controller plasmids pCRB DRT7VTPLux*500CFP and pCRB DRT7VTPLux*500YFP lacking either the bicistronic operon P_Las81*_-*T7RNAP[1-179]*-*mVenus* or P_Lux76*_-*T7RNAP[180-880]*-*mTurquoise2*, respectively, were constructed. Each half-circuit controller plasmid was introduced into *TOP10 E. coli* alongside the RFP reporter plasmid p4g3R. CFP, YFP, and RFP fluorescence intensities were measured over time as a function of 3OC6HSL and 3OC12HSL concentrations using a plate fluorometer-based assay (see Methods for details). As expected, the half-circuit bicistronic controller plasmid pCRB DRT7VTPLux*500CFP responded to increasing concentrations of 3OC6HSL by induction of corrected CFP fluorescence intensity, the half-circuit bicistronic controller plasmid pCRB DRT7VTPLux*500YFP to increasing concentrations of 3OC12HSL by induction of corrected YFP fluorescence intensity (Tab. 1, Supplementary Fig. S4a-b). Induction of corrected RFP intensity from the reporter plasmid was negligible for both half-circuit bicistronic controller plasmids. Under the same assay conditions, the complete bicistronic controller plasmid pCRB DRT7VTPLux*500 produced substantial levels of corrected RFP fluorescence intensity if exposed to both 3OC6HSL and 3OC12HSL. Notably, we observed a reduction in corrected YFP (up to 29 ± 7%) and CFP (up to 46 ± 9%) fluorescence intensities under conditions of high RFP induction.

**Table 1.** Comparison of bicistronic controller plasmid pCRB DRT7VTPLux*500 and its constitutive half circuits. *TOP10 E. coli* were co-transformed by an *mRFP1*-expressing reporter of T7RNAP activity p4g3R and the bicistronic controller plasmid pCRB DRT7VTPLux*500 or one of its constitutive half-circuits. By contrast to pCRB DRT7VTPLux*500, pCRB DRT7VTPLux*500CFP and pCRB DRT7VTPLux*500YFP lacked either the bicistronic operon P_Las81*_-*T7RNAP[1-179]*-*mVenus* or P_Lux76*_-*T7RNAP[180-880]*-*mTurquoise2*, respectively. The behavior of bicistronic controller plasmid variants was tested alongside the RFP reporter plasmid under a two-dimensional titration of 3OC6HSL and 3OC12HSL using a plate fluorometer-based assay (see Methods for details). Reported for each construct is maximum observed fluorescence intensity, corrected for background signal present in absence of externally supplied homoserine lactones. Shown is the s.d. of average values yielded between 3 biological replicate experiments performed on different days. Activity plots are shown in Supplementary Fig. S4. Individual induction curves are shown in Supplementary Fig. S5.

| Max corrected fluorescence intensity [RFU] |  |  |  |
| --- | --- | --- | --- |
|  | CFP | YFP | RFP |
| pCRB DRT7VTPLux*500CFP | 7,000 ± 1,000 | 9 ± 9 | 100 ± 20 |
| pCRB DRT7VTPLux*500YFP | 1,200 ± 200 | 5,000 ± 1,000 | 150 ± 20 |
| pCRB DRT7VTPLux*500 | 5,000 ± 1,000 | 5,000 ± 1,000 | 11,000 ± 3,000 |

While measurement of corrected RFP intensity under two-dimensional titration of homoserine lactones showed AND gate-like behavior of our circuit (Supplementary Fig. S4c), we sought to further reduce background induction in the absence of 3OC12HSL. Corrected RFP fluorescence intensity in the absence of 3OC6HSL was already low. We replaced RBS_B0033_ (relative strength 500 arbitrary units) regulating expression of *T7RNAP[1-179]* in pCRB DRT7VTPLux*500 by a series of weaker ribosomal binding sites (relative strength 250, 100, and 50 arbitrary units) designed by the Ribosome Binding Site Calculator^43^. Comparing the resulting bicistronic controller plasmid variants using plate fluorometry, we found pCRB DRT7VTPLux*250 to most sensitively respond to increasing concentrations of 3OC12HSL (Supplementary Fig. S6). In a two-dimensional titration of 3OC6HSL and 3OC12HSL, the improved three-color circuit composed of bicistronic controller pCRB DRT7VTPLux*250 and reporter plasmid p4g3R induced corrected RFP fluorescence intensity to over 8-fold of the maximum background detected in the absence of either input signal (Supplementary Fig. S8). The improved synthetic three-colour circuit correctly exhibited the behavior of a synthetic transcriptional AND gate responsive to two different homoserine lactones (Fig. 4).

**Figure 4.**
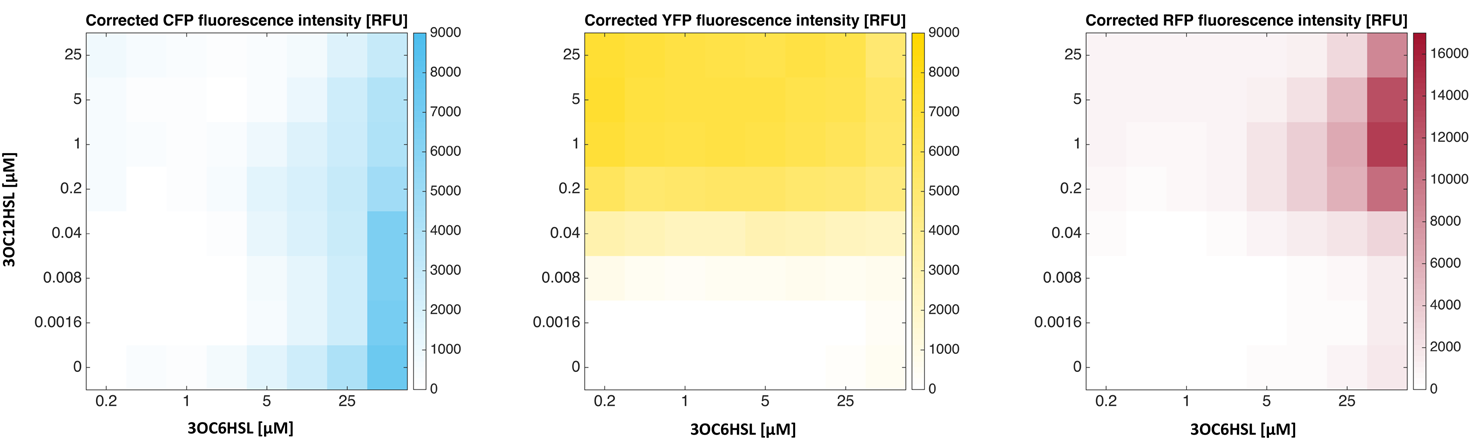
An improved synthetic three-color circuit implementing AND gate-like behavior. The behavior of the high-dynamic-range bicistronic controller plasmid pCRB DRT7VTPLux*250 was tested alongside the RFP reporter plasmid p4g3R under a twodimensional titration of 3OC6HSL and 3OC12HSL using a plate fluorometer-based assay (see Methods for details. Reported for each condition is fluorescence intensity corrected for background signal present in absence of externally supplied homoserine lactones. Plots show average values from 3 biological replicate experiments performed on different days. Corresponding s.d. values are shown alongside individual induction curves in Supplementary Fig. S8.

### Homoserine lactone-mediated patterning of bacterial populations

To test whether our three-color synthetic transcriptional AND gate was responding to homoserine lactones in surface culture as well as in suspension, we employed a previously reported spatial assay based on membranes printed with hydrophobic grids^24^. Bacterial populations are confined to quadrants with well-defined geometry, separated from one another while allowing homoserine lactone-mediated communication across the substrate. Following inoculation of membranes with dilute bacterial cultures, changes in CFP, YFP, and RFP fluorescence intensities in response to environmental signals can be monitored across the grid over time using a custom macroscopic imaging system (see Methods for details).

First, we uniformly inoculated 64 membrane quadrants with bacteria co-transformed by the RFP reporter plasmid p4g3R and the improved bicistronic controller plasmid pCRB DRT7VTPLux*250. In this experiment, membranes were placed on minimal nutrient agar containing (i) no added homoserine lactones, (ii) 25 μM 3OC6HSL, (iii) 1 μM 3OC12HSL, or (iv) both of the previously mentioned. The concentrations of signalling molecules were chosen to match the condition of highest induction of RFP fluorescence intensity observed in suspension culture (see Fig. 4). Solid culture assays performed at 37°C as outlined above confirmed that presence of both homoserine lactones was required for substantial induction of corrected RFP fluorescence intensity, suggesting AND gate-like behaviour (Fig. 5a). Corrected CFP and YFP fluorescence intensities were reduced to background level under the condition of co-induction compared to presence of either signalling molecule alone.

**Figure 5.**
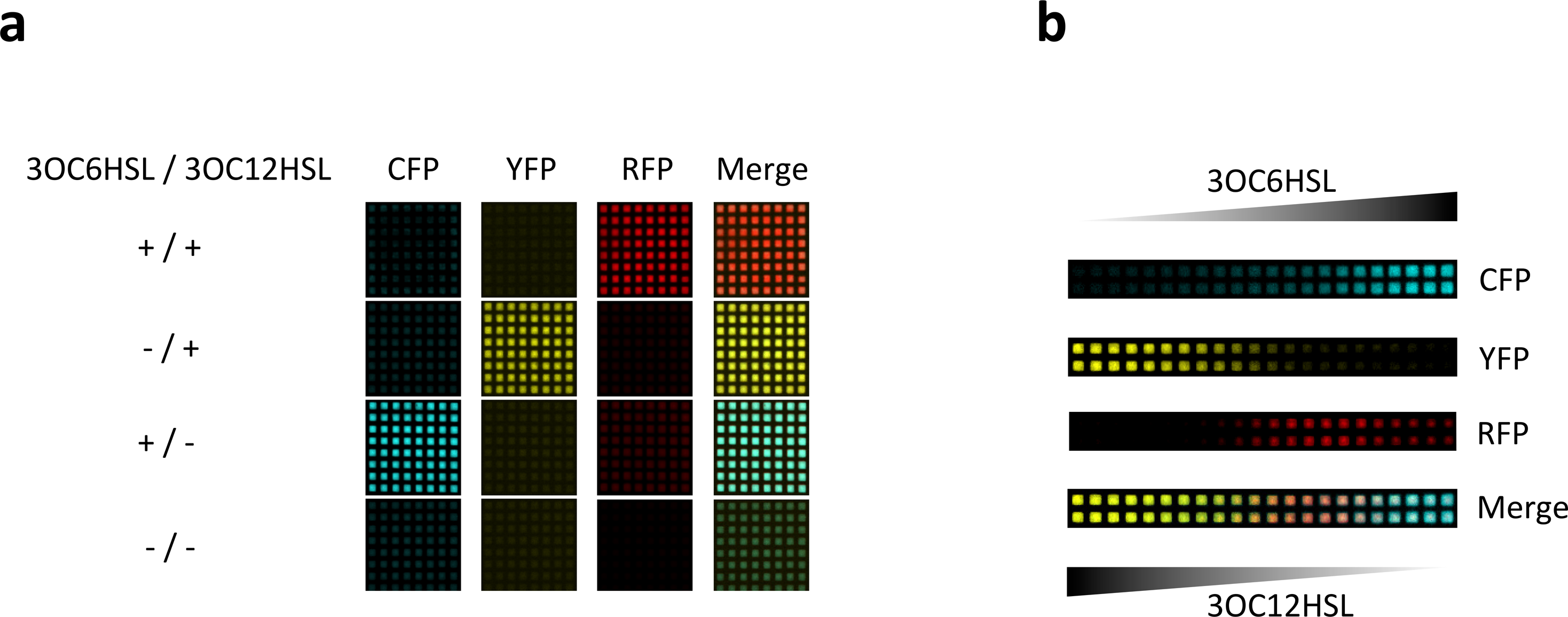
Surface-based patterning of bacterial gene expression by means of a three-color AND-gate in response to externally supplied homoserine lactones. (**a**) Surface-based circuit behavior in response to homoserine lactones present in the growth medium at uniform concentration. *TOP10 E. coli* co-transformed by the RFP reporter plasmid p4g3R and the improved bicistronic controller plasmid pCRB DRT7VTPLux*250 (see Fig. 3) were incubated on membranes printed with hydrophobic ink, placed on minimal agar containing different combinations of 3OC6HSL (25 μM) and 3OC12HSL (1 μM). Images shown were captured at *t* = 1,500 min (time relative to start of incubation). Corresponding corrected fluorescence intensities are shown in Supplementary Fig. S9. (**b**) Surface-based AND gate behavior in response to homoserine lactone gradients. *TOP10 E. coli* cells co-transformed by controller and reporter plasmids as above were incubated on membranes placed on minimal agar lacking supplemented homoserine lactones. Instead, aqueous solutions containing 500 μM 3OC6HSL or 200 μM 3OC12HSL, respectively, were spotted next to the cells on either side and left to diffuse into the bacterial population from opposite directions. Images shown were captured at *t* = 3,000 min (time relative to start of incubation). Corrected fluorescence intensities recorded over time are shown in Supplementary Fig. S10.

Next, we sought to test whether the same synthetic three-color circuit could implement surface-based spatial patterning across a bacterial population in response to gradients of diffusing homoserine lactones. For this purpose, we inoculated rows of quadrants with double-transformed cells on agar lacking supplemented homoserine lactones. At a distance of one quadrant to either end of the inoculated rows, we then spotted solutions of 3OC6HSL (500 μM) or 3OC12HSL (200 μM), and allowed the inducers to diffuse into the bacterial population from opposite directions. Over a course of 50 hours, we observed the emergence of a three-color pattern across the bacterial population: with fluorescence of CFP and YFP mirroring the opposite gradients of 3OC6HSL and 3OC12HSL, RFP fluorescence was found to peak near the centre of the inoculated rows where the two gradients overlap (Fig. 5b).

### Programmed emergence of a gene expression domain in the absence of externally applied inducer gradients

Finally, we implemented a greater degree of autonomy in our synthetic three-color patterning system by enabling bacterial cells to produce and secrete their own inducers. The improved bicistronic controller plasmid pCRB DRT7VTPLux*250 and the RFP reporter p4g3R had been characterized both in suspension (see Fig. 4) and in surface culture (see Fig. 5). We introduced a third plasmid into *E. coli*: the sender plasmids pSB1C3 I0500 (LuxI/LasI) embraced either *luxI* (encoding an enzyme producing 3OC6HSL; Stevens & Greenberg 1997) or *lasI* (encoding an enzyme producing 3OC12HSL; Schuster et al. 2004) under control of the P_BAD/araC_ promoter system.

Utilizing solid culture assays, we inoculated membrane grids with two adjacent populations of the different triple-transformed cell types: each population harboured (i) the RFP reporter, (ii) the improved bicistronic controller, and (iii) either the LuxI or the LasI sender plasmid. In the absence of arabinose, fluorescence intensity was low in all three channels. In the presence of 25 mM arabinose, corrected CFP and YFP fluorescence intensities were substantially induced in cells expressing LuxI and LasI, respectively (Fig. 6). Using HSL standard curves prepared under the same conditions (Supplementary Fig. S12), the effective concentrations of 3OC6HSL and 3OC12HSL in their respective domains of synthesis were roughly estimated at 0.2-1 μM and 1 μM, respectively, at quadrants farthest from the interface. A symmetrical domain of high RFP activity spontaneously emerged at the interface of 3OC6HSL- and 3OC12HSL-sending cell populations. Our synthetic three-color patterning circuit generated programmed emergence of this new gene expression domain in absence of externally applied signalling gradients, implementing an autonomous mechanism for hierarchical patterning.

**Figure 6.**
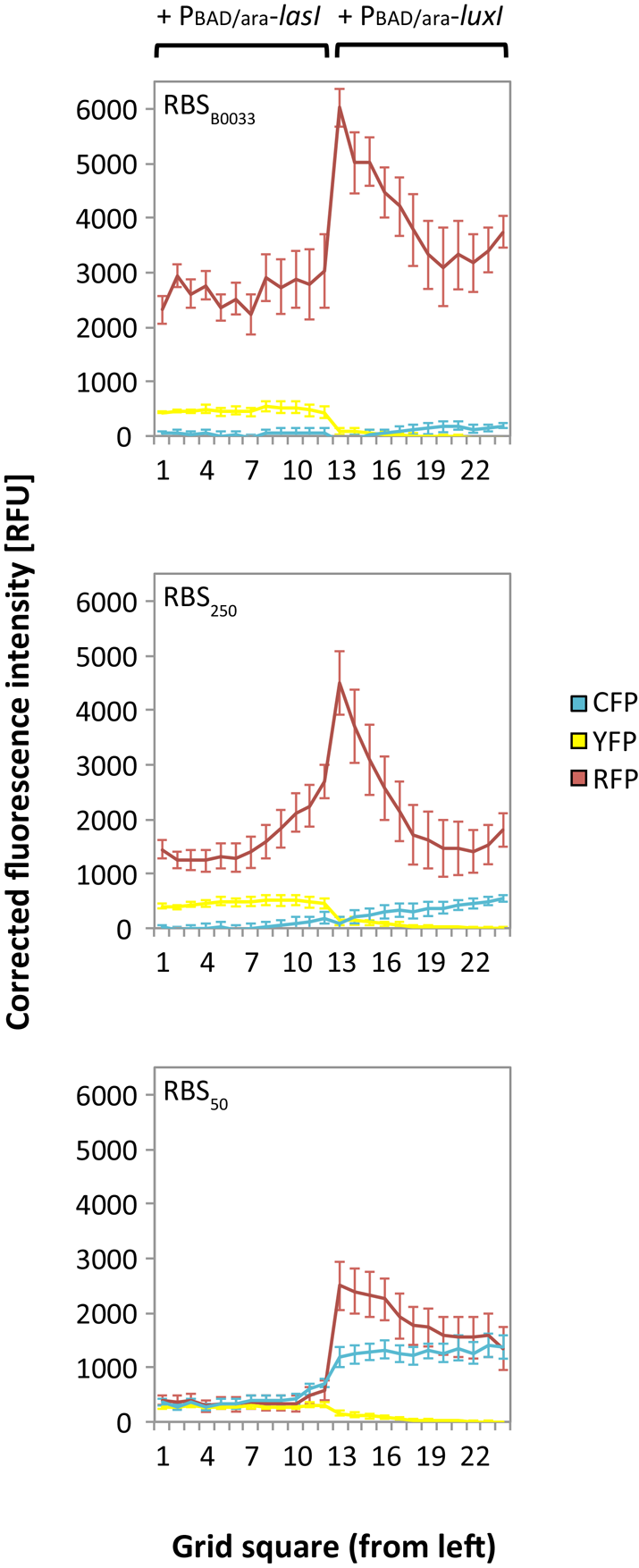
Programmed emergence of an RFP-expressing domain across bacterial populations in the absence of externally applied homoserine lactones. *TOP10 E. coli* were co-transformed by the RFP reporter plasmid p4g3R, a bicistronic controller plasmid pCRB DRT7VTPLux*(500/250/50), and a sender plasmid pSB1C3 I0500 (LuxI/LasI) encoding either *luxI* or *lasI* under control of the arabinose-inducible P_BAD/araC_ promoter. Adjacent populations of the different triple-transformed cell types were incubated on membranes placed on minimal nutrient agar which has or has not been supplemented with 25 mM arabinose. Reported for each set of quadrants equidistant from the genotype boundary (12 each) is fluorescence intensity corrected for background signal present in absence of arabinose, at *t* = 3,000 min (time relative to start of incubation). Error bars represent the s.d. of average values between the 12 equidistant quadrants on each membrane. Corresponding images captured at *t* = 3,000 min are shown in Supplementary Fig. S11.

To assess the robustness of this behavior, we subjected the suboptimal bicistronic controller plasmids pCRB DRT7VTPLux*500 and pCRB DRT7VTPLux*50 (see Supplementary Fig. S6) to the same solid culture experiment. Compared to RBS_250_ in pCRB DRT7VTPLux*250, presence of the stronger RBS_B0033_ regulating expression of *T7RNAP[1-179]* in pCRB DRT7VTPLux*500 slightly increased corrected YFP fluorescence intensity while reducing corrected CFP fluorescence intensity to approximately a third of its original value. pCRB DRT7VTPLux*500 exceeded pCRB DRT7VTPLux*250 by a third in peak corrected RFP fluorescence intensity while background activity in the *luxI*-expressing domain more than doubled, as assessed by quadrants farthest from the interface.

Compared to RBS_250_ in pCRB DRT7VTPLux*250, the weaker RBS_50_ regulating expression of *T7RNAP[1-179]* in pCRB DRT7VTPLux*50 decreased corrected YFP fluorescence intensity by over a third while more than doubling CFP fluorescence intensity. Peak corrected RFP fluorescence intensity was reduced almost to half of its original value, with high background activity across the *luxI*-expressing domain. These results marked the relative performance of bicistronic controller plasmid variants in plate fluorometer-based assays (see Supplementary Fig. S6) predictive of their suitability to implement surface-based hierarchical patterning, and underlined the importance of empirical circuit optimization.

## Discussion

Hierarchical induction of gene expression domains is a key mechanism in living organisms relying on patterning for establishment of their physical shape^12,14,44^. However, this mechanism has received little attention from synthetic biologists in efforts to emulate developmental processes fundamental to morphogenesis^5–11,24^. In this work, we implemented programmed spatial patterning resembling hierarchical induction in populations of bacterial cells. Our approach was based on a synthetic three-color genetic circuit embracing a split T7RNAP. This enzyme was controlled by two different signalling molecules derived from bacterial quorum sensing (3OC6HSL and 3OC12HSL)^16–18^. The homoserine lactones shaped a first layer of patterning and spatially controlled expression of split T7RNAP which then established a second layer of patterning by transcriptional activity from its cognate T7 promoter.

In developing our synthetic circuit, we first sought to identify a variant of split T7RNAP exhibiting the highest dynamic range among a range of candidates. To this end, we extended a previously reported ratiometric strategy for robust quantification of promoter activity^30,31^ to T7RNAP-driven gene expression: in our ratiometric reporter plasmids, the T7 promoter controlled expression of *mVenus* YFP^32^ as primary output of T7RNAP activity. Simultaneous monitoring of constitutive *mTurquoise2* CFP^33^ expression from the same plasmid allows correction of the primary output for the overall gene expression capacity of the cell under given environmental conditions.

Comparing different ratiometric reporter plasmids for T7RNAP activity, we found that a variant combining a weak mutant T7 promoter P_T7_(-3G)^37^ with a low copy number origin of replication pSC101^45^ exhibited the highest dynamic range (see Fig. 2a). This observation reflects the well-established fact that high cellular activity of T7RNAP can result in toxicity^46^, likely due to decoupling of T7RNAP-driven transcription and host translation^47–49^ and/or depletion of cellular resources such as nucleotides or amino acids^50^. Indeed, the mutation R632S, thought to reduce the processivity of T7RNAP, has been shown to alleviate toxicity associated with this expression system in *E. coli*^39^. In our ratiometric assays, presence of the R632S mutation in T7RNAP increased the maximum induction from the T7 promoter by almost 4-fold (see Fig. 2b). Splitting T7RNAP between residues 179 and 180^25^ is another modification to the enzyme which has been shown to both reduce its processivity^51^ and to alleviate toxicity in *E. coli*^26^. In our experiments, (i) introducing the R632S mutation into intact T7RNAP and (ii) splitting the enzyme increased the maximum induction of activity from the T7 promoter to a similar extent. The inducer concentration required for half maximal activity from the T7 promoter was also similarly increased in T7*RNAP and split T7RNAP (~30-fold) compared to T7RNAP (see Supplementary Fig. S2). Our results confirm an earlier report of split T7RNAP resolving loss of activity from the T7 promoter upon high enzyme induction^26^.

We also tested variants of split T7RNAP modified by either the R632S mutation, the addition of SynZIP protein-protein interaction domains^40 41^, or both of these features. Interestingly, with split T7RNAP, the individual modifications decreased the maximum induction of activity from the T7 promoter to approximately a third of its original value, with combination of the R632S mutation and SynZIP domains reducing it another 10-fold. The inducer concentration required for half maximal activity was also markedly increased for these variants (over 100-fold compared to T7RNAP; see Supplementary Fig. S2). We concluded that splitting T7RNAP sufficiently reduced its processivity to alleviate toxic effects and further modification of this variant unnecessarily compromised its activity.

As the next step towards a synthetic patterning circuit, we sought to build on the proven capacity of split T7RNAP to implement AND-logic^26–28^, and to make this behavior dependent on spatial coincidence of the diffusible signalling molecules 3OC6HSL and 3OC12HSL. In our bicistronic controller plasmid pCRB DRT7VTPLux*500 (see Fig. 3), the N- and C-terminal fragments of split T7RNAP were controlled by promoters induced by either of the two homoserine lactones. Those promoters were derived from our previously reported system for orthogonal intercellular signalling^24^, but modified by mutations shown to reduce basal activity from the Lux promoter by approximately 6-fold^42^. As visual indicators of expression of the individual T7RNAP fragments *in vivo*, we chose to include the fluorescent reporter genes *mTurquoise2* (downstream of the 3OC6HSL-responsive P_Lux76*_ promoter and *T7RNAP[180-880]*) and *mVenus* (downstream of the 3OC12HSL-responsive P_Las81*_ promoter and *T7RNAP[1-179]*) in the bicistronic controller plasmid. Both were correctly induced by their cognate homoserine lactones in suspension (see Tab. 1, Supplementary Fig. S4). However, a notable level of activity from the T7 promoter as indicated by the RFP reporter plasmid was observed for the bicistronic controller system in the absence of supplemented 3OC12HSL (up to approximately a third of maximum circuit induction). This suggested that the two constitutive half circuits were not perfectly orthogonal, but the promoter P_Las81*_ was induced by 3OC6HSL to some extent. To alleviate this effect, we tested several weak ribosomal binding sites designed by the Ribosome Binding Site Calculator^43^ to control expression of *T7RNAP[1-179]* downstream of P_Las81*_ (see Supplementary Fig. S6). We identified a variant RBS_250_ capable of enhancing the maximum circuit induction in presence of both homoserine lactones to over 8-fold of the maximum background detected in the absence of either input signal (see Fig. 4, Supplementary Fig. S8). Interestingly, we observed that corrected fluorescence intensity in both the CFP and the YFP channels was reduced by over 25% under conditions exhibiting high levels of RFP fluorescence compared to conditions inducing either controller half circuit alone. This may be a consequence of rapid transcription of the *mRFP1* gene by T7RNAP relative to *mVenus* and *mTurquoise2* transcribed by endogenous *E. coli* RNA polymerase^46–49^, leading to the latter two transgenes being partially out-competed for cellular gene expression capacity. Posttranscriptional limitation of fluorescent protein expression is further suggested by the inverse correlation between CFP expression and RBS strength controlling the YFP operon during application of bicistronic controller plasmids in surface culture (see Fig. 6). As an alternative explanation for the reduction in fluorescence intensities observed, the possibility of quenching of *mTurquoise2* and *mVenus* in presence of *mRFP1* cannot be excluded.

Our synthetic three-color patterning circuit also behaved in an AND gate-like manner in surface culture (see Fig. 5), albeit with a reduced dynamic range (see Supplementary Fig. S9). The previously mentioned reduction in CFP and YFP fluorescence intensities under conditions of high RFP expression was more pronounced in the surface-based experiment than in suspension culture. One possible explanation for this observation lies in comparatively slow bacterial growth rates during surface culture^52^, which can lead to intracellular accumulation of proteins^53^: in the case of T7RNAP components, protein build-up may promote leaky expression of *mRFP1* from the T7 promoter and increase out-competition of *mVenus* and *mTurquoise2* for cellular gene expression capacity, as discussed above. Intracellular accumulation of fluorescent proteins may also promote quenching due to molecular crowding effects.

Both in suspension (see Fig. 4) and on minimal nutrient agar containing a uniform concentration of homoserine lactones (see Fig. 5a, Supplementary Fig. S9), we observed induction of RFP fluorescence to be primarily limited by 3OC12HSL. This is consistent with low-level response of P_Las81*_ to 3OC6HSL. In an earlier report, we have shown fluorescent proteins directly controlled by P_Lux76_ and P_Las81_ to be induced by their cognate homoserine lactones 3OC6HSL and 3OC12HSL in an effectively orthogonal manner^24^. In this work, T7RNAP serves as a potent amplifier of primary promoter activity, magnifying a previously undetectable level of crosstalk.

Despite this effect, our synthetic three-color circuit was capable of generating the emergence of a new pattern of gene expression in response to opposite inducer gradients applied externally (see Fig. 5b) or produced autonomously (see Fig. 6). Under the former condition, we observed a comparatively flat peak in corrected RFP fluorescence across the isogenic bacterial population, likely shaped by the high concentration of externally applied signalling molecules. By contrast, a sharp domain strongly expressing RFP was generated at the interface of 3OC6HSL- and 3OC12HSL-sending cell populations.

This demonstration opens a door to programming hierarchical patterning behavior across populations of cells. For example, replacing the *mRFP1* gene in our synthetic three-color circuit by a biosynthetic enzyme producing a third diffusible signal orthogonal to 3OC6HSL and 3OC12HSL would allow programmed patterning of a multicellular population into 5 different domains across 3 hierarchical levels (Fig. 7). Candidate orthogonal signalling systems for this purpose have already been described ^22,23,54^. In principle, the population-based AND gate can be iterated with *n* orthogonal signalling systems to generate (2 × n) − 1 different domains.

**Figure 7.**
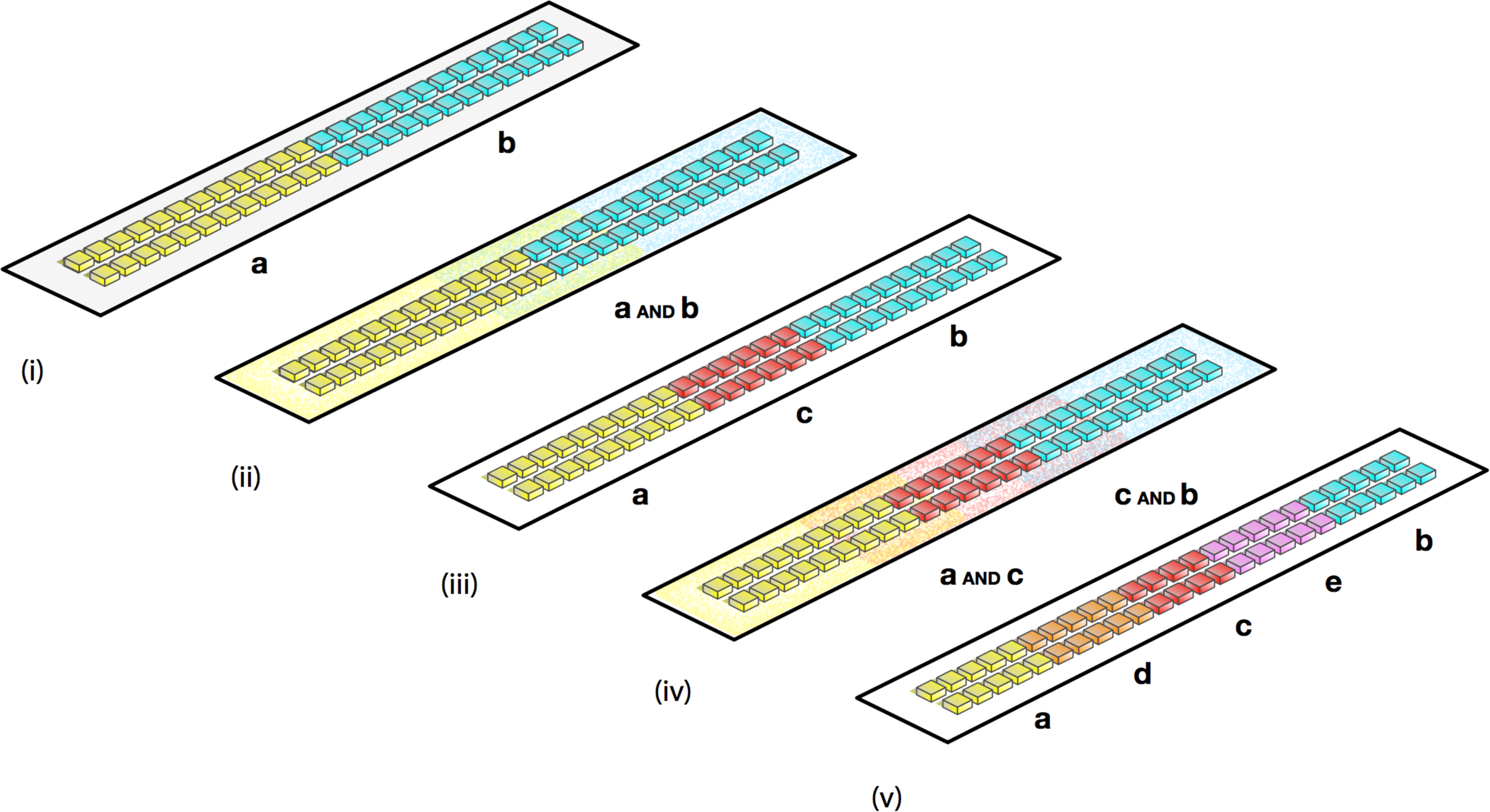
A general framework for hierarchical patterning of bacterial populations. (i) An initial asymmetry splits a bacterial population into two different gene expression domains (yellow and cyan). The yellow domain produces a diffusible morphogen a, the cyan domain a diffusible morphogen b. (ii) Morphogens a and b coincide at the boundary of the yellow and cyan domains. (iii) Population-based logic induces a red gene expression domain in response to a AND b. The red domain produces an additional diffusible morphogen c. (iv) Morphogens a and c coincide at the boundary of the yellow and red domains, morphogens c and b at the boundary of red and cyan domains. (v) Population-based logic induces an orange gene expression domain in response to a AND c, and a purple gene expression domain in response to c AND b. The orange and purple domains produce additional diffusible morphogens d and e, respectively. This scheme may be iterated to generate (2 × *n*) − 1 different domains with *n* orthogonal signalling systems.

The implementation of higher-order hierarchical patterning processes may become limited by insufficient signal-to-noise ratio in effector expression. In this respect, our approach to implementation of two-layer synthetic hierarchical patterning underlines the importance of empirically characterizing individual genetic components prior to their application in a complex synthetic circuit. As illustrated by the performance of suboptimal bicistronic controller plasmid variants in solid culture assay (see Fig. 6), modulation of a single ribosomal binding site was capable of qualitatively changing circuit behavior: exchange of RBS_250_ for the stronger RBS_B0033_ or the weaker RBS_50_ upstream of *T7RNAP[1-179]* both shifted the balance between CFP and YFP expression in the first layer of patterning and affected the signal-to-noise ratio in RFP expression in the second layer of patterning. Our success in designing a synthetic genetic circuit implementing the desired behavior was dependent on empirical selection of a high-dynamic-range variant of split T7RNAP through ratiometric assays (see Fig. 2) and on translational tuning of the resulting AND gate through plate fluorometer assays (see Supplementary Fig. S6).

In nature, morphogen gradients mediate developmental patterning at a high level of precision in face of various perturbations, supported by sophisticated genetic circuits and layers of feedback mechanisms^55–57^. Likewise, the robustness of synthetic hierarchical patterning could be increased by supplementary mechanisms implementing mutual inhibition, positive feedback, or morphogen degradation. In terms of the circuit introduced herein, mutual inhibition between domains at the first layer of patterning could be implemented by additional expression of *lasR* under control of P_Lux76*_, and of *luxR* under control of P_Las81*_. Positive feedback could be implemented by expression of a *T7RNAP*&gene under control of the T7 promoter, rendering maintenance of the second layer of patterning independent from the first layer inducers 3OC6HSL and 3OC12HSL. To alleviate potential propagation of signalling crosstalk to higher levels of patterning, the first layer inducers 3OC6HSL and 3OC12HSL could further be specifically degraded at the second layer of patterning. This could be achieved by expression of a quorum quenching enzyme - such as the lactonase *aiiA*^58^ - under control of the T7 promoter. Feedback mechanisms like the ones outlined can be used to consolidate induced domains and to promote correct transmission of spatial information across layers of complexity. As a test-bed for the development of new synthetic feedback circuits, our system for synthetic hierarchical patterning promises to facilitate future efforts at creating custom multicellular organisms for applications in bioproduction, remediation, or medicine.

## Methods

### Plasmid construction

All plasmids (listed in Supplementary Tab. S16) were constructed using Gibson assembly^59^ with parts obtained from the MIT Registry of Standard Biological Parts (http://partsregistry.org), from Addgene (www.addgene.org), or synthesized by Integrated DNA Technologies (Coralville, IA, USA), and are available on Addgene. Identities and source of backbone vectors, genes, and regulatory elements used in this work are summarized in Supplementary Tab. S13-S15. Sequences are available on Genbank (accession numbers KY643824 and KX986152 to KX986173). Controller devices encoding intact and split variants of *T7RNAP* were based on plasmids acquired from Shis & Bennett^26^. Bicistronic controller devices were based on a double receiver plasmid previously reported^24^. All cloning was performed in *TOP10 E. coli* (Invitrogen, Waltham, MA). With exception of ratiometric reporter plasmids for T7RNAP activity (*T7 Express E. coli*; New England Biolabs, Ipswich, MA, USA) all analysis was also carried out in *TOP10 E. coli*.

### Plate fluorometer assays

Plate fluorometer assays were conducted as previously described^24^, with minor modifications: overnight cultures were diluted 1:100 in M9 medium supplemented with 0.2%w/v casamino acids, 0.4%w/v glucose, and inducers to the concentrations described, and loaded in a final volume of 200 μL per well onto a clear-bottom 96-well microplate (Greiner, Kremsmünster, Austria). Measurements of CFP (excitation 430/10 nm, emission 480/10 nm), YFP (excitation 500/10 nm, emission 530/10 nm), and RFP (excitation 550/10 nm, emission 610/20 nm) fluorescence intensities, and OD_600_ were taken approximately every 12 min for 100 cycles (approximately 19 hrs) in a BMG FLUOstar Omega plate fluorometer (BMG Labtech, Ortenberg, Germany) at 37 °C, under shaking at 200 rpm between readings. Data analysis was performed using the Genetic Engineering of Cells (GEC) modeling and design environment (version 6.12.2014) as previously described^30^.

### Solid culture assays

Solid culture assays were performed as previously reported^24^, with minor modifications: single colonies were picked from LB agar plates and grown overnight in supplemented M9 medium with appropriate antibiotics (50 μg/mL carbenicillin and 50 μg/mL kanamycin for double-transformed cells; 50 μg/mL carbenicillin, 50 μg/mL kanamycin, and 25 μg/mL chloramphenicol for triple-transformed cells). Cultures were diluted 1:100, then grown into exponential phase to an optical density at 600 nm of 0.3. This dilute culture was spotted onto Iso-Grid membranes (40 × 40 quadrants of 1 mm per side; Neogen, Lansing, MI, USA) placed on 1.5%_w/v_ agar plates containing the same supplemented M9 growth medium. The culture was plated at a volume of 0.5 μL per quadrant. Plates were incubated and imaged in a custom imaging device which has been described in detail elsewhere^24^. For quantification of fluorescence intensities, mean pixel grey values localized to individual Iso-Grid quadrants across the CFP, YFP, and RFP channels were extrapolated using the open-source Fiji distribution of ImageJ^60^.

### Data availability

The authors declare that all data supporting the findings of this study are available within the paper and its Supplementary Information files or are available from the corresponding author on request.

## Acknowledgements

C.R.B. acknowledges support from the Gates Cambridge Trust. P.K.G. acknowledges support from the John Templeton Foundation (Grant No. 15619: “Mind, Mechanism and Mathematics: Turing Centenary Research Project”). J.H. acknowledges support from the Biotechnology and Biological Sciences Research Council and the Engineering and Physical Sciences Research Council (OpenPlant Grant No. BB/L014130/1) and EC FP7 project no. 612146 (PLASWIRES).

## Author contributions

C.R.B., P.G. and J.H. conceived and contributed key ideas to the development of the project; C.R.B. and P.G. designed and performed the experiments; C.R.B. analysed the data; J.H. supervised the project. All authors contributed to the manuscript.

## Additional information

**Accession codes:** The sequences of 23 plasmids used in this study have been submitted to the GenBank nucleotide database under accession codes KY643824 and KX986152 to KX986173.

**Supplementary Information** accompanies this paper

**Competing financial interests:** The authors declare no competing financial interests.

